# Global control of cellular physiology by biomolecular condensates through modulation of electrochemical equilibria

**DOI:** 10.1101/2023.10.19.563018

**Authors:** Yifan Dai, Zhengqing Zhou, Kyeri Kim, Nelson Rivera, Javid Mohammed, Heileen Hsu-Kim, Ashutosh Chilkoti, Lingchong You

## Abstract

Control of the electrochemical environment in living cells is typically attributed to ion channels. Here we show that the formation of biomolecular condensates can modulate the electrochemical environment in cells, which affects processes globally within the cell and interactions of the cell with its environment. Condensate formation results in the depletion or enrichment of certain ions, generating intracellular ion gradients. These gradients directly affect the electrochemical properties of a cell, including the cytoplasmic pH and hyperpolarization of the membrane potential. The modulation of the electrochemical equilibria between the intra- and extra-cellular environments by biomolecular condensates governs charge-dependent uptake of small molecules by cells, and thereby directly influences bacterial survival under antibiotic stress. The shift of the intracellular electrochemical equilibria by condensate formation also drives a global change of the gene expression profile. The control of the cytoplasmic environment by condensates is correlated with their volume fraction, which can be highly variable between cells due to the stochastic nature of gene expression at the single cell level. Thus, condensate formation can amplify cell-cell variability of the environmental effects induced by the shift of cellular electrochemical equilibria. Our work reveals new biochemical functions of condensates, which extend beyond the biomolecules driving and participating in condensate formation, and uncovers a new role of biomolecular condensates in cellular regulation.

---

Biomolecular condensates are ubiquitous meso-micro scale cellular structures that regulate various cellular processes, such as transcription, stress response, and RNA splicing (*1, 2*). The formation of condensates is driven by the phase transition of multivalent associative biomacromolecules (*3*). Studies have uncovered the molecular signatures that encode the phase behaviors in biomacromolecules and the mechanisms by which cells use compartmentalization of specific biomacromolecules to modulate cellular functions (*4–6*). Besides the participation of biomacromolecules in condensate formation, experimental and theoretical studies have discovered that biomolecular condensates can also spatially segregate ions (*7–12*), which can result in a pH gradient, and thereby establish an electric potential gradient between the dilute and the dense phases (*7, 8*). A prime example of such an ion gradient is the pH difference between the nucleolar compartment and the nucleoplasm (*13*).

Small ions are the most abundant cellular constituent other than water (*14*). The spatial distribution of ions dictates the cellular electrochemical potential, which serves as a fundamental driving force of biochemical reactions and cellular signaling (*15*), such as the proton motive forces set by a membrane electric potential (*16*). Our present understanding of the mechanism by which ions are modulated intracellularly primarily originates from studies of ion channels in cellular membranes (*17*).

Simulations based on liquid-state theory predict that complex coacervation, a type of phase separation, can result in an electrostatic potential difference—a Galvani potential—between the dilute and the dense phase (*8–10, 18, 19*). Indeed, our recent study demonstrated that condensate formation by certain proteins (i.e., resilin-like-polypeptides) could generate an electrical potential gradient between the dilute and the dense phases (*7*), and thereby create an interfacial electrical double layer that can drive redox reactions. These findings imply a possibility that the phase transition of biomacromolecules can affect the electrochemical equilibria of the system. However, it is unclear whether condensate formation can modulate the electrochemical environment in the cytoplasm, thereby affecting the intracellular electrochemical equilibrium, and whether this would have an impact on cellular physiology and function.

Here, we demonstrate the role of condensates in modulating the cytoplasmic ion environment and the functional consequences of this condensate-mediated shift in the electrochemical equilibria of cells. We find that the selective partitioning of ions in a cell by condensate formation can alter the cytoplasmic ionic environment in terms of its spatial pH and the distribution of specific ions. The asymmetry in the distribution of specific ions in the cytoplasm modulates the cellular membrane potential. We further show that these features have a long-reaching impact, including modulation of the interactions between the cell and its environment and regulation of gene expression on a global scale in cells. Moreover, the capacity for this global control depends on the volume fraction of condensates within a cell, which generates highly variable cellular behaviors due to the stochasticity in gene expression. This heterogeneity in cellular behaviors due to condensate formation is in sharp contrast to the role of condensates in mediating the protein concentration homeostasis of the phase separating protein, as shown previously (*20*). This work uncovers a new functional role for condensates in the regulation of electrochemical equilibria and a new mechanism by which condensates can modulate cellular function.

## Results

### Condensation formation modulates cytoplasmic ion abundance

We previously found that condensates (the dense phase) possess a distinct pH environment from the cytoplasmic environment (the dilute phase) (*7*) (**Fig. 1A**). To understand the effects of condensates on cytoplasmic ion environment, we first expanded these results by using the same synthetic intrinsically disordered protein (synIDP) (*21*), consisting of a resilin-like-polypeptide (RLP; amino acid sequence: Ser-Lys-Gly-Pro-[Gly-Arg-Gly-Asp-Ser-Pro-Tyr-Ser]_20_-Gly-Tyr), which undergoes phase separation via an upper critical solution temperature (UCST) phase transition (*22, 23*). This synIDP shares similar sequence features and the same thermodynamic driving force as many native IDPs that can drive condensate formation, such as FUS and hnRNP A1 (*4, 24*). We placed the synIDP in a pET-24 plasmid under the control of a T7 promoter containing a *lac* operator, which is inducible by isopropyl ß-D-1-thiogalactopyranoside (IPTG) in *E. coli* cells expressing LacI (*23*).

**Fig. 1.**
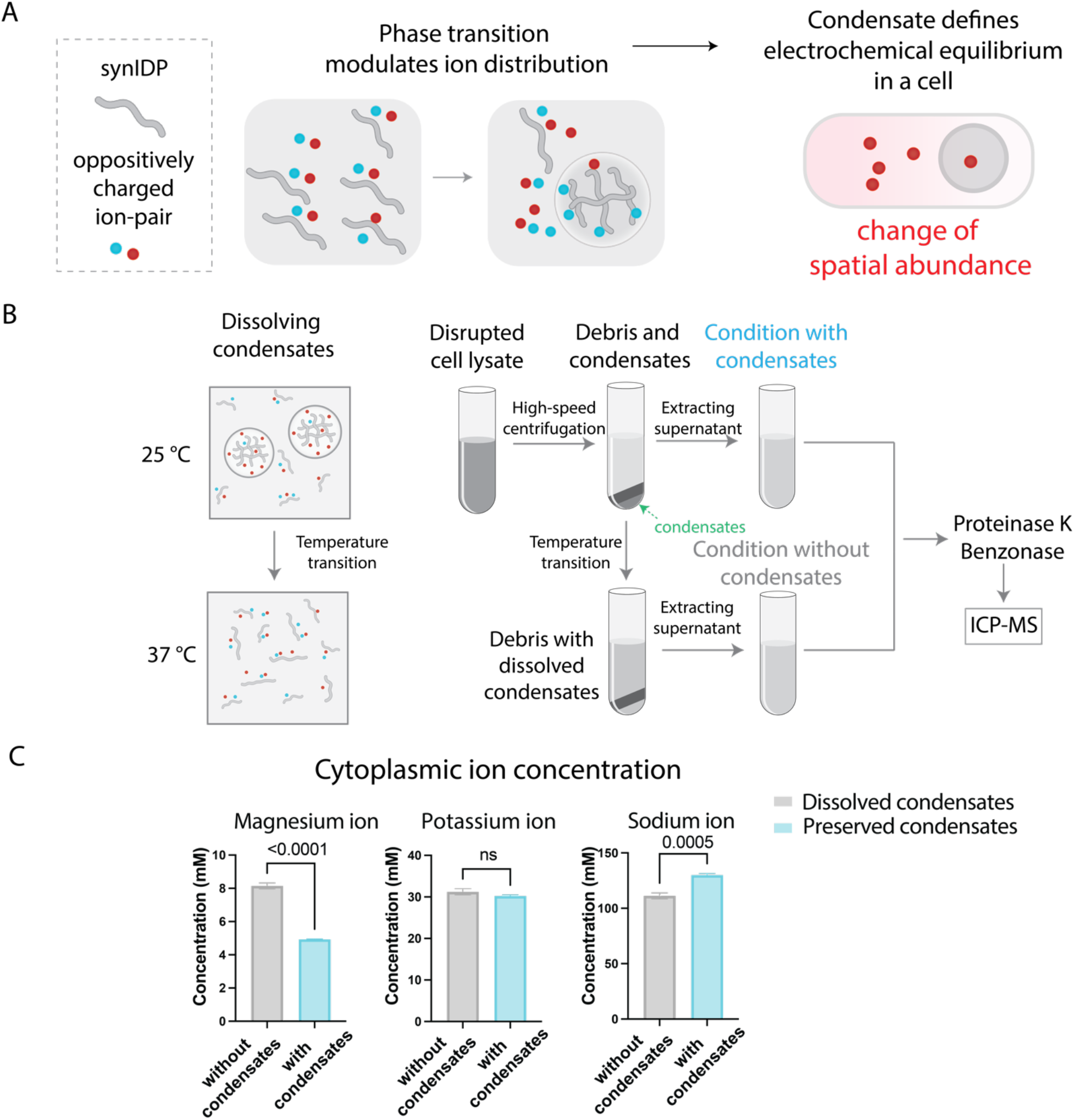
Condensate formation modulates cytoplasmic ion abundance. **A,** Phase transition of associative macromolecules can lead to segregation or exclusion of specific types of ions, which can modulate the intracellular ion distribution. **B,** Inductively coupled plasma mass spectrometry (ICP-MS) analysis of the effects of phase transition of an RLP on cytoplasmic ion concentration. Upper critical solution temperature (UCST) transition behavior is utilized to dissolve RLP condensates by increasing the solution temperature above the transition temperature of the RLP. The supernatant fraction corresponds to the condition of cytoplasm. The supernatant is treated with proteinase K (a final concentration of 0.4 units/mL) and benzonase nuclease (25 units/mL) before analysis by ICP-MS. The ICP-MS result was converted back to estimated cellular ion concentration through cell density and dilution ratio based on the sample processing methods. **C,** Cytoplasmic ion concentration of magnesium, potassium and sodium ions at conditions with or without RLP condensates. Two-tailed t-test for statistical analysis. ns = non-significant.

Using a pH-dependent ratiometric dye, C-SNARF-4-AM(*25*), we found that condensate formation by the RLP led to a more acidic cytoplasmic pH and a more alkaline environment in the condensate (**fig. s1a, b**) within a wide range of the external pH conditions (**fig. s1c**). We next investigated condensates formed by an elastin-like polypeptide (ELP) with a sequence of [Val-Pro-Gly-Val-Gly]_40_ that can undergo phase transition by a lower critical solution temperature (LCST) transition, but cannot generate an ion gradient (*7*). We found that the ELP condensates did not significantly affect the cytoplasmic pH (**fig. s2**).

The pH gradient in the cell arises from the enforcement of charge neutrality in each phase under the equilibrium of electrochemical potential (*9, 10*). This implies that the difference in pH between the two phases would either be the cause or the result of a selective partitioning of other types of small ions (e.g., sodium) in the two phases (*26–28*). To identify these ions and measure their partitioning, we used inductively coupled plasma mass spectrometry (ICP-MS) (*29, 30*). We used a previously established protocol to isolate condensates from mechanically disrupted cells and exploited the temperature sensitivity of the UCST phase behavior of the RLP to dissolve the separated dense phase after isolation (*22*) (**Fig. 1B**). This strategy was applied to generate samples with and without condensates. We then measured the abundance of ions in the samples with and without condensates by ICP-MS and calculated the ion concentrations in the condensate and the cytoplasm.

We measured the concentration of three different cations, sodium (Na^+^), magnesium (Mg^++^), and potassium (K^+^) ions (**Fig. 1C**), as their distributions are known to control the intracellular pH and various cellular processes, such as ribosome activity and stress resistance (*31–35*). We found that condensates reduced the cytoplasmic concentration of Mg^++^ by ∼2-fold. As the typical volume ratio between condensates and cytoplasm is ∼1:4, this implies that condensates have a 5-fold higher Mg^++^ concentration relative to the cytoplasm. In contrast, Na^+^ was excluded from condensates, with condensate formation increasing the cytoplasmic Na^+^ concentration by 19% compared with that of samples without condensates. The exclusion of Na^+^ by the condensates might be due to the change of the hydration property of Na^+^ ions in the condensate solvent environment (*36*). We did not observe an impact of condensate formation on the distribution of K^+^ between the two phases. We next measured the ion distribution in the case of ELP condensates. We found that formation of ELP condensates did not lead to a statistically significant change in the concentration of any of these ions in the cytoplasm (**fig. s3**).

These observations confirm the correlation between a pH gradient between the dilute and dense phases and the capability of condensates to modulate the flux of small ions. The asymmetric distribution of ions between the dilute and the dense phases further suggests that condensate formation can modulate the electrochemical equilibria within a cell.

### Volume fraction of condensates modulates cytoplasmic ion and generates noise in cells

During phase separation, thermodynamic constraints dictate that for a one-component phase separating system(*37*), such as the RLP system, phase separation provides a mechanism to reduce noise in the cytoplasmic concentration of the phase separating component (*20*). For instance, as the concentration of the RLP in the cytoplasm exceeds its C_sat_, the system enters the two-phase region of the phase diagram and the concentration in each phase stays approximately the same, whereas the increase in the total amount of RLP in a cell is reflected by an increase in the volume fraction of the RLP condensates (*38, 39*) (**fig. S4a**).

Thus, if RLP expression is variable between cells due to stochastic gene expression, this variability will be reflected in the volume fraction of condensates of individual cells (*40*). To test this notion, we modeled the dynamics of phase separation coupled with stochastic gene expression by adopting the framework of Kolesin et al. (*20*). Our model accounts for both intrinsic noise due to small numbers of interacting molecules and extrinsic noise in the transcriptional rate (**Supplementary Note 1**)(*41*). Consistent with a previous study (*20*), simulations show that condensate formation led to reduced variability (measured by the coefficient of variation, or CV) in the cytoplasmic concentration of the condensate-forming protein, relative to that of mRNA and total protein (**Fig. 2A**). In contrast, the volume fraction of the condensates in a cell had similar variability as the total concentration of mRNA of a cell (**Fig. 2B**).

**Fig. 2.**
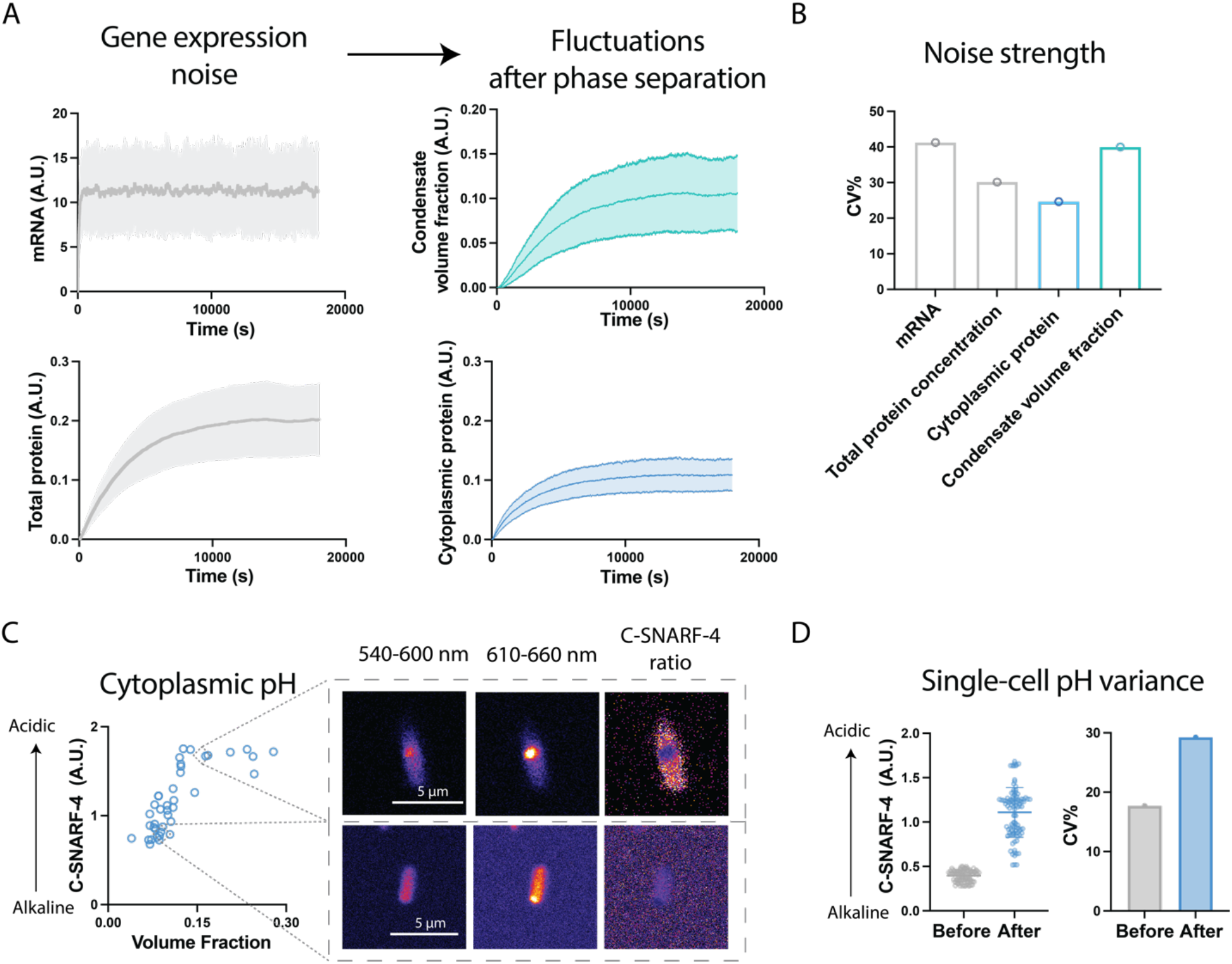
Condensate formation increases the heterogeneity of cellular physiology. **A,** The mean and variance of gene expression and the mean and variance of condensate volume fraction and cytoplasmic protein from a stochastic model of gene expression. **B,** Amplitude of noise in the total mRNA concentration, total protein concentration, cytoplasmic protein (dilute phase) concentration and condensate volume fraction as a function of expression time in the model. **C,** Dependency of cytoplasmic pH on condensate volume fraction from cells induced for expression for different amounts of times (30 min, 60 min, 180 min and 300 min). Each data point represents a single cell. **D,** Single-cell pH variance before and after condensate formation. Cells were analyzed at different time points before (30 min, 0 min before induction) and after condensate formation (60 min, 120 min and 180 min after induction). Coefficient of variance (CV) = (standard deviation/mean)*100. N = 75 individual cells.

As the total capacity of a phase to partition or exclude ions should be proportional to the total volume of the phase, we hypothesized that the heterogeneity in the volume fraction of condensates between cells would drive a corresponding heterogeneity in the cytoplasmic ion environment between individual cells. To test this notion, we first measured the cytoplasmic pH of individual cells at different time points after induction of RLP expression that corresponded to different condensate volume fractions. We found that a larger condensate volumetric fraction corresponds to a more acidic cytoplasmic pH value, as seen by a higher C-SNARF-4 signal (**Fig. 2C**). Moreover, cells with condensates showed a 1.5-fold increase in the variability of their cytoplasmic pH compared to cells without condensates (**Fig. 2D & fig. S5a**).

To verify whether the modulation of the cytoplasmic chemical environment is due to the fluctuations of the condensate volume fraction instead of the fluctuations of the cytoplasmic RLP concentration, we measured the cytoplasmic protein concentration before and after condensate formation in individual cells using a previously established fluorescence based method by fusing the RLP with a monomeric green fluorescent protein (mEGFP) (*23*). We found a slightly reduced variability of cytoplasmic protein concentration after condensate formation (**fig. s5a&b**), consistent with our model prediction and previous experimental observation (*20*). Thus, the amplified cytoplasmic pH variability across a cell population was not due to the fluctuation of protein concentration in the cytoplasm. Taken together, our results confirm that for the environmental effect brought about by condensate formation, phase separation amplifies the cell-cell variability in the intracellular chemical environment.

### Intracellular phase separation modulates membrane potential

Cells typically exert control over their intracellular ion content through ion channels (*42–44*), which in turn modulate the membrane potential (*45*). As membrane potential is highly sensitive to the intracellular electrochemical environment(*45–48*), with the new equilibrium state of intracellular ion distribution established by condensates, we hypothesized that condensate formation can modulate the electrochemical equilibrium between extracellular and intracellular ions and thereby change the membrane potential (**Fig. 3A**).

**Fig 3.**
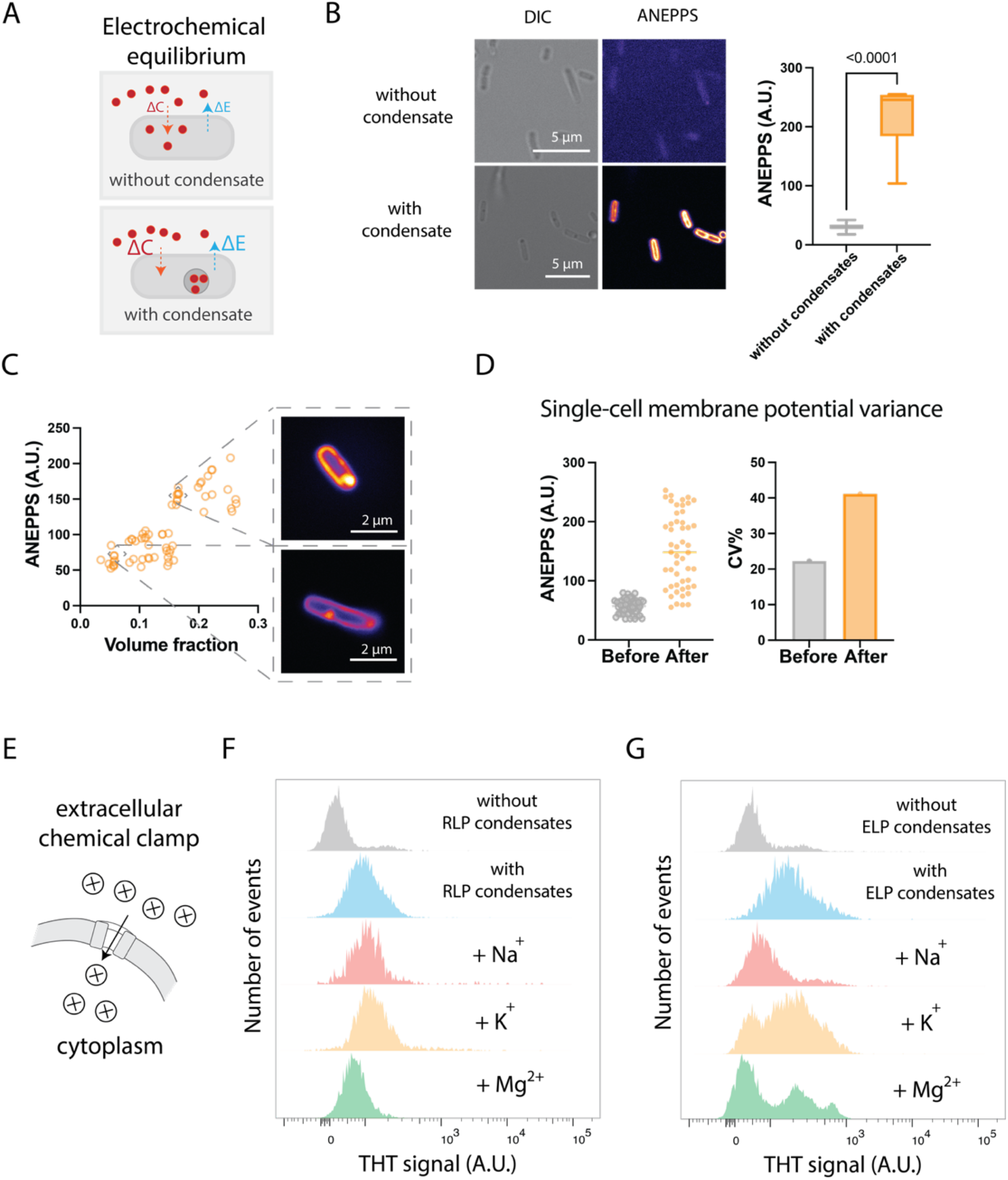
Biomolecular condensates regulate membrane potential, establishing a new electrochemical equilibrium between the intracellular and extracellular environment. **A,** Electrochemical potential equilibrium between extracellular and intracellular environments is modulated by condensate formation. **B,** Sample images of DI-4-ANEPPS dye that measures the change of membrane potential upon condensate formation. A higher fluorescence corresponds to membrane hyperpolarization. Two-tailed t-test for statistical analysis. **C,** Dependency of membrane potential on condensate volume fraction of cells induced for RLP expression for different amounts of times (30 min, 60 min, 180 min and 300 min). Each data point represents a single cell. **D,** Single-cell membrane potential variance before and after condensate formation. Cells were analyzed at different time points before (45 min, 0 min before induction) and after condensate formation (60 min, 180 min and 240 min after induction). Coefficient of variance (CV) = (standard deviation/mean)*100. N = 50 individual cells. **E,** Ion supplementation as chemical clamp to depolarize the membrane potential for identification of ion types that drive membrane potential hyperpolarization. **F,** Flow cytometry analysis of ThT signal of single cells based on different conditions of cells without or with RLP condensates and supplemented with different extracellular ion sources (NaCl, KCl and MgCl_2_). 14.3 % ThT positive for without condensates condition; 72.4% ThT positive for with condensates condition; 78.3% ThT positive for with condensates condition supplemented with NaCl; 90.0% ThT positive for with condensates condition supplemented with KCl; 50.1% ThT positive for with condensates condition supplemented with MgCl_2_. **G,** Flow cytometry analysis of ThT signal of single cells based on different conditions of cells without or with ELP condensates and supplemented with different extracellular ion sources (NaCl, KCl and MgCl_2_). 21.0 % ThT positive for without condensates condition; 83.4% ThT positive for with condensates condition; 40.3% ThT positive for with condensates condition supplemented with NaCl; 73.3% ThT positive for with condensates condition supplemented with KCl; 44.7% ThT positive for with condensates condition supplemented with MgCl_2_.

To test this hypothesis, we measured the membrane potential of cells with and without RLP condensates by incubating log-phase growing cells in Hank’s balanced salt solution (HBSS) at the same cellular density. We first used an Amino Naphthyl Ethenyl Pyridinium dye (DI-4-ANEPPS) to evaluate the change of membrane potential; upon binding to the cellular membrane, the dye generates a fluorescence signal whose intensity increases with the hyperpolarization of the membrane potential based on a local electric field-dependent change in the electronic structure of the dye (*49, 50*). Indeed, the fluorescence intensity of membrane-associated ANEPPS was ∼7.5-fold higher in cells containing condensates than those without condensates (**Fig. 3B**). The condensates also showed a substantial increase in the ANEPPS signal compared to the cytoplasm, which is consistent with our previous study that the interface of condensates has an electric potential gradient (*7*). In single cells where condensate formation was induced for different durations of time, the membrane potential increased with the volume fraction of condensates (**Fig. 3C**). Moreover, condensate formation led to a CV% of 40 % of membrane potential signal (**Fig. 3D**), which is similar to the variability of volume fraction (**Fig. 2B**). These results demonstrate that the capability of condensates to modulate intracellular electrochemical equilibria can affect membrane potential, providing another biophysical basis of the global impact of condensates on cellular physiology.

The change of membrane potential may have two distinct causes: 1) the distribution of charged species in the extracellular and intracellular environment is different before and after condensate formation, and 2) the function of the ion channel is specifically inhibited due to condensate formation. To explore the possible driving forces, we used a fluorescence-based thioflavin-T (ThT) assay (*46*), the partitioning of which into the cells is based on the Nernst potential (*51*), thereby serving as an orthogonal strategy to profile the membrane potential based on the intracellular abundance of ThT. We combined this approach with a chemical-clamp method (*46*) to evaluate the ion drivers that mediate the change of membrane potential (**Fig. 3E**). We observed that cells with RLP condensates exhibited an 80-fold increase in the ThT fluorescence signal than those without condensates (**Fig. 3F**), indicating a more negative membrane potential (a state of hyperpolarization) upon condensate formation (*46, 52*). This phenomenon triggered by condensates is similar to the passive effect on the membrane potential if there is an efflux of ions (*53, 54*), thereby confirming that the change of membrane potential arises from the capacity of condensates to modulate the distribution of ions between the intracellular and extracellular milieu. It is noteworthy that the ThT fluorescence signal of cells with condensates had a broader distribution compared to the ThT signal of cells without condensates, again confirming the amplified heterogeneity in the membrane potential due to condensate formation.

To identify the ion species that modulate the membrane potential, we supplemented the test buffer (HBSS with ThT dye) with different monovalent and multivalent cations (Na^+^, K^+^, Mg^++^) that are known to affect membrane potential (*50*) and measured the change of the ThT signal for cells with condensates. Only the addition of Mg^++^ substantially repressed the ThT fluorescence with a decrease in ThT positive cells from 72.4% to 50.1 % (**Fig. 3F**), restoring the membrane potential to a level similar to that of cells without condensates. This observation indicates that the hyperpolarized membrane potential is mediated through the partitioning of Mg^++^ into the condensates, consistent with the ion distribution measurement by ICP-MS (**Fig. 1E**).

Interestingly, ELP condensates also caused a change in the cell membrane potential as measured by the ThT assay (**Fig. 3G**). Using the same chemical-clamp approach, we found that the supplementation of sodium ion mediates the strongest repression of membrane potential with a decrease in ThT positive cells from 83.4% to 40.3 %. Though we did not find ELP condensates could strongly partition ions (**fig. s3**), the change of membrane potential upon the formation of ELP condensates is still significant compared to cells without ELP condensates. As the ELP phase separates through a LCST mechanism (*55*), in which the phase transition of the ELP leads to the change of the water content in the dense phase, it thereby modulates the water abundance in the cytoplasm. We speculated that it is possible that this change in the water content of the cytoplasm results a change of the osmotic equilibrium between the intracellular and extracellular environment (*56*), which is known to influence the membrane potential (*57, 58*). These observations show that condensates, due to their ability to affect the chemical environment of the cells, can strongly regulate the electrochemical equilibrium between intracellular and extracellular environment.

We next explored whether the ability of condensates to modulate the membrane potential can be programmed at the protein sequence level. We reasoned that by changing the overall charge of the phase separating proteins, their condensates would exhibit different degrees of ion partitioning, thereby showing a different capacity to perturb the membrane potential. As a proof-of-principle experiment, we fused the same RLP sequence with super folder green fluorescence protein (sfGFP), a negatively charged fluorescent protein, which imparts a net 6 negative charges (**fig. s6**). We found that the RLP-sfGFP fusion decreased the ANEPPS signal by 3.5-fold compared to the RLP alone, suggesting that the condensate-mediated electrochemical equilibrium is potentially programmable at the sequence level of a condensate forming protein.

### Condensate modulates membrane permeability and antibiotic tolerance

A change in the membrane potential could have a profound impact on small molecule signaling (*50, 59–61*), by tuning the direction and magnitude of the flux of small molecules (*45, 46, 48, 52*). We hypothesized that this modulation can potentially affect the survival of cells exposed to functional small molecules (*62–64*), such as antibiotics, based on their electrostatic charge (**Fig. 4A**). To test this hypothesis, we measured the growth recovery of back-diluted log-phase cells with and without RLP condensates in M9 media containing different antibiotics. In all media conditions, cells with or without condensates exhibited different growth dynamics (**Fig. 4B, C & D)**. In the absence of antibiotics, cells with condensates showed a 2.4-fold longer lag time than cells without condensates (**Fig. 4B**), suggesting a burden caused by condensate formation on the recovery of cells. However, when exposed to negatively charged antibiotics (namely ampicillin and carbenicillin) at sublethal concentrations, cells with condensates either showed a faster recovery from the lag phase or a higher maximum growth rate (**Fig. 4C, E & F**). This increased tolerance to ampicillin and carbenicillin likely resulted from the reduced uptake of these negatively charged antibiotics due to the condensate-mediated hyperpolarized membrane potential.

**Fig 4.**
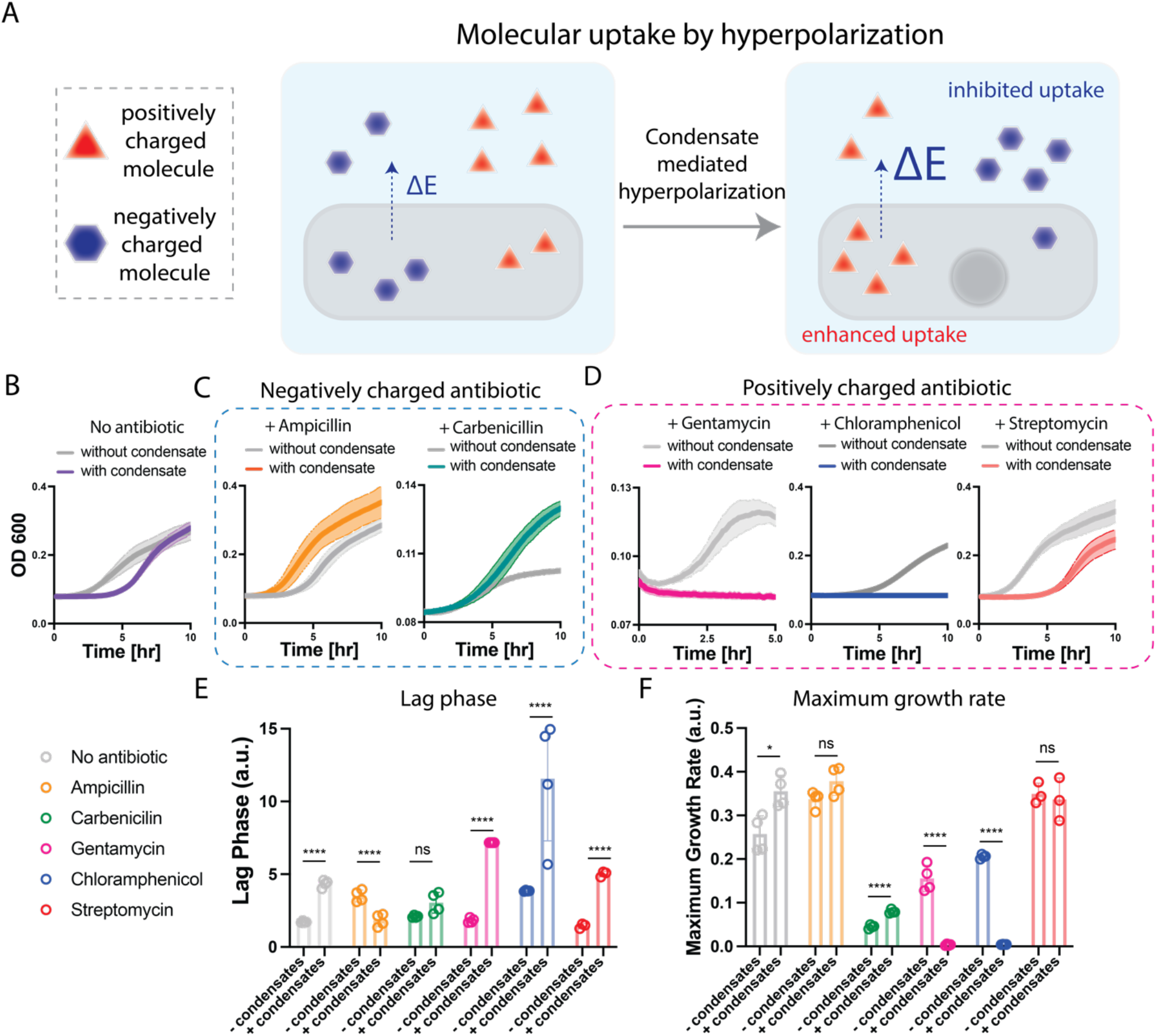
Biomolecular condensates dictate membrane molecular uptake and cellular fitness. **A,** Condensates mediate hyperpolarization of cellular membrane, thereby changing the uptake of molecules based on their types of electrostatic charge. **B,** Growth of cells with or without condensates in M9 minimum medium. **C,** Growth of cells with or without condensates in M9 minimum medium with negatively charged antibiotics, including 2 µg/mL ampicillin (left panel) and 1 µg/mL carbenicillin (right panel). **D,** Growth of cells with or without condensates in M9 minimum medium with positively charged antibiotics. 1 µg/mL gentamicin (left panel), 6 µg/mL chloramphenicol (middle panel) and 2 µg/mL streptomycin (right panel). **E,** Comparison of the extracted growth lag phase of cells with or without condensates under the treatment of different antibiotics. ****, P<0.0001; ns, non-significance; based on unpaired t-test. **F,** Comparison of the extracted maximum growth rate of cells with or without condensates under the treatment with different antibiotics. *, P = 0.0117; ****, P < 0.0001; ns, non-significance; based on unpaired t-test.

In contrast, condensate formation greatly amplified the impact of positively charged antibiotics (namely gentamycin, chloramphenicol, and streptomycin). When applied at sublethal concentrations (**Fig. 4 D, E & F**), all three positively charged antibiotics significantly inhibited the recovery of cells containing condensates, which showed at least a 3-fold longer lag phase than the cells without condensates. Cells with condensates treated with gentamycin and chloramphenicol did not grow at all. The enhanced susceptibility likely resulted from increased uptake of these positively charged antibiotics due to the condensate-mediated hyperpolarized membrane potential.

The mechanism that condensate-mediated membrane hyperpolarization can direct antibiotic uptakes provides an intuitive explanation of a previous discovery that a native condensate formed by HslU protein could enhance bacterial tolerance to antibiotics including ampicillin and carbenicillin (*64*). It is likely that condensate formation of HslU led to hyperpolarization of the membrane potential, thus suppressing the uptake of negatively charged antibiotics and thereby enhancing cell survival. If so, we would expect condensate formation by HslU would change the membrane potential and enhance the efficacy of positively charged antibiotics.

To test this hypothesis, we expressed the HslU protein from a pET 24 plasmid in *E. Coli*. Using the ThT assay, we found that HslU condensate formation led to an increase in membrane potential (**fig. s7a**), like RLP condensates. Consistent with the previous study (*64*), we found that cells with condensates showed increased tolerance to a negatively charged antibiotic, ampicillin (**fig. s7b**) (*64*). In contrast, consistent with our prediction, comparing the growth recovery response of cells with and without HslU condensates, cells with HslU condensates were more susceptible to a positively charged antibiotic, gentamycin, when applied at sublethal concentrations, compared to the growth response of cells without condensates (**fig. s6c**).

### The electrochemical features of condensates affect cellular processes

The preceding evidence shows that the formation of biomolecular condensates can modulate the global electrochemical equilibria in cells. We reasoned that this change in electrochemical equilibria can, in turn, drive a global change in gene expression in cells. To test this notion, we prepared cells with and without condensates and measured their global gene expression using RNA-sequencing (**Fig. 5a**).

**Fig 5.**
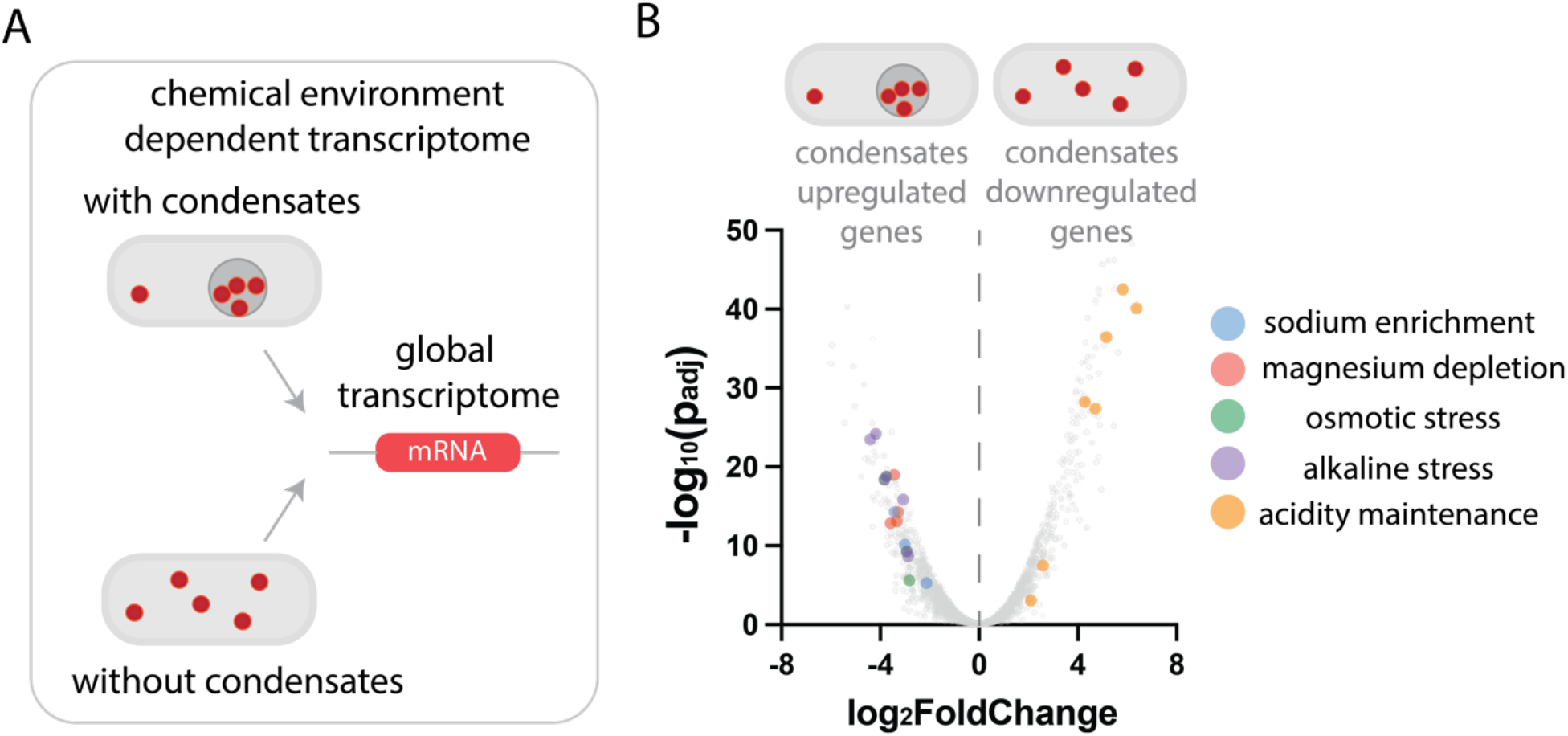
Condensate dependent change of global transcriptome. **A,** Cells with or without RLP condensates were subjected to RNA-sequencing analysis to evaluate differential gene expression profiles between samples. **B,** Volcano plot (fold change of mRNA level between samples vs. adjusted p-value) shows the distribution of transcriptomes in cells with and without condensates. Statistically significant and featured transcripts are color coded based on the chemical features upregulating the downstream gene expression.

1557 genes were differentially expressed between cells with and cells without condensates with an adjusted P-value < 0.0499. (**Fig. 5b**, see Supplementary table for gene ID and count values). Of these, we assessed the expression profiles of genes that can be regulated by the changes of cellular chemical environment, including pathways regulated by ion-dependent transcription factors, osmotic stress and pH. Compared with cells without condensates, cells with condensates upregulated the expression of all genes in Mg^++^ transporter pathways (*mgt, mgr, bor* gene clusters), which are known to be activated when Mg^++^ is limiting (*65*), as well as three of four genes under control of the Na^+^-dependent transcription factor NhaR (*66, 67*). A set of genes involved in osmotic stress response (e.g., *rrrD, psp, ybd, bet, aqp* gene clusters and stress response sigma factor *rpoE*) (*68–71*) were also upregulated in cells with condensates. These altered gene expression profiles align with the selective enrichment of Na^+^ and depletion of Mg^++^ in the cytoplasm upon condensate formation (**Fig. 1**). In addition, several genes involved in pH homeostasis, such as alkaline stress genes and acidity maintenance genes (e.g., slp, asr, glsA, ybaT, Adi clusters) (*72*) were also differentially expressed between cells with condensates and cells without condensates.

These data highlight two critical points. First, our targeted analysis of gene expression profiles aligns with the observed environmental effects of condensate formation, such as the intracellular ion distribution. Second, these data suggest that the function of condensates goes well beyond the specific set of biomolecules that participate in condensate formation, as shown by the finding that condensates can mediate global changes in gene expression profiles within the cell.

## Discussion

The generation of ion gradients by associative polymers that undergo phase separation has been studied using liquid-state theory-based simulations of the complex coacervation of asymmetric mixtures of polyelectrolytes (*8, 18*). The asymmetry in charge distribution brought about by macromolecules in one phase needs to be balanced by ions to realize the electrochemical potential equilibrium that is dictated by phase equilibrium (*26*). However, the extent to which electrochemical equilibria in living cells can be modulated by condensates, and how that affects cellular physiology, is unclear. This study provides direct evidence of how the phase separation of associative biomacromolecules can establish an ion gradient within cells. We demonstrate that the ion gradient drastically affects the electrochemical equilibria of a cell by modulating: 1) intracellular ion homeostasis, as seen by the pH/ion concentration difference between the cytoplasm and condensates; and 2) the membrane potential, by inducing hyperpolarization of the cell membrane.

Intracellular ion homeostasis is critical to cellular physiology (*59, 65, 73*), as the precise control of ion concentrations and their spatial distribution is the foundation of biochemical activities within a cell (*74*), including transcription (*75*), translation (*76*), antibiotic resistance (*46*). Our results show that cells can utilize condensate formation as an intracellular mechanism to regulate spatial distribution of specific ions within a cell, which provides a mechanism to regulate cellular electrochemical equilibria that is orthogonal and complementary to ion channels.

Are these features of condensates likely to generally hold true across diverse condensates? We suggest that this is likely based on the conserved molecular interactions that drive phase separation of diverse condensate-forming IDPs and high net charge nature of typical components that participate in condensate formation, such as nucleic acids (*77*). From the perspective of biomacromolecules that undergo phase transitions, such as IDPs and nucleic acids (*77, 78*), a key difference before and after phase transition is the switch of the interactions between with the aqueous solvent and with themselves. Before they undergo a phase transition, biomacromolecules are solvated by water molecules with exposed charged residues that are screened by counterions. After phase transition, the biomacromolecules percolate with each other, thereby releasing counterions and water molecules. Due to the limited flexibility of the backbone of IDPs and nucleic acids (*9, 79*), a complete screening of charged residues through intra- and inter-molecular interactions is likely impossible. Therefore, the requirement of charge neutrality within a phase requires that additional ions would enter the condensate from the cytoplasm to screen the excess charge. This generates an asymmetry in the ion distribution between phases. Furthermore, this asymmetry in ion distribution can also be possibly triggered by interactions that do not involve charge-based electrostatic interactions. For example, the distinct solvent environment in the dense phase as a result of a change in the abundance of water content or micropolarity (*80, 81*) can direct the partial solvation of ions, which in turn can create a distinct ion environment in the dense phase, thereby promoting the formation of an ion gradient. Hence, all these confounding factors suggest that biomolecular condensates are an asymmetric mixture of components, which suggests a possibly generalizable feature of condensates in mediating cellular electrochemical equilibria and establishing an electrochemical gradient within a cell.

From a cellular standpoint, the diffusion length scale of ions is much longer than macromolecules (*14*), so that the chemical effect exerted by condensates can dictate long-range cellular control as evidenced by the change in the interactions of cells with their environment. Specifically, the intracellular ion gradient established by condensates is rebalanced with the extracellular environment through ion channels in the cellular membrane, which leads to a change of the membrane potential (*45, 46, 53*). In turn, these changes affect intracellular gene expression on a global scale and the response of cells to environmental perturbations (Fig. 4). These results clearly demonstrate that condensates can encode cellular functions beyond the specific functions that are encoded in the biomolecules that form the condensates.

A final notable finding of this study relates to the effect of stochasticity of gene expression at the single cell level. Gene expression—including that of IDPs that form the condensate—is stochastic at the single cell level. Though phase separation can reduce the concentration fluctuation of the phase separating component in the cytoplasm (*20*), the cell-cell variability in the total concentration of the IDP is manifested by a similar cell-cell variability in the volume fraction of the condensate. The cell-cell variability in the volume of the condensate then translates to similar cell-cell variability in the effect of the cytoplasmic ion environment in the cell and the extent of hyperpolarization of the cell membrane.

In summary, our finding uncovers a new biochemical function of condensates in modulating electrochemical equilibria in cells, and which in turn affects global cellular physiology. This function has a distinct effect in generating cellular noise, which further suggests that the cooperativity and signaling between different condensates and cellular components may be a critical—and as yet unstudied— phenomenon by which the interplay of different biomolecular condensates in cells controls cellular homeostasis. This work uncovers a new level of the functional complexity of biomolecular condensates, and expands our understanding of their cellular functions beyond those encoded in the biomolecules that participate in condensate formation.

## ACKNOWLEDGMENTS

We thank the Duke Light Microscopy Core facility for experimental support. This work was supported by the Air Force Office of Scientific Research (FA9550-20-1-0241) to L.Y. and A.C. and by the NIH (R35-GM127042) to A.C. We appreciate discussions with and comments from Rohit V. Pappu in the Department of Biomedical Engineering at Washington University in St. Louis, Clifford Brangwynne in the Department of Chemical and Biological Engineering at Princeton University, and Amy Gladfelter and Christine Roden in the Department of Cell Biology at Duke University.

## AUTHOR CONTRIBUTIONS

Y.D. conceived of the idea and Y.D., L.Y. and A.C. devised the study. Y.D., Z.Z., K.K., N.R., and J.M. performed the experiments. Y.D., L.Y., A.C. wrote and revised multiple versions of the manuscript. All the authors read and contributed revisions. All the authors analyzed the data and contributed to discussions. A.C. and L.Y. acquired funding.

