## Supplementary Materials for "Global control of cellular physiology by biomolecular condensates through modulation of electrochemical equilibria"

This PDF includes:

- Supplementary Text 1
- Materials and Methods
- Figure S1-S8
- Table S1-2

**Supplementary Text 1.** Gene expression noise leads to condensate volume heterogeneity.

Change in the volume fraction of the condensate within the cell could drive the global shift in the chemical environment in cytoplasm, thus gene expression patterns. Here we show the generic condensate-forming protein expression noise as one of the potential causes of condensate volume fraction heterogeneity.

We adopted the model by Klosin et al. (*1*) describing gene expression and phase separation. Here we enclose a brief description of the model. The model consists of a generic gene expression model (*2*) and a free-energy-governed stochastic protein exchange process between protein in dilute and dense phases.

$$\emptyset \xrightarrow{\Lambda_r} \mathrm{RNA} \xrightarrow{n_r/\tau_r} \emptyset$$

$$\emptyset \xrightarrow{\lambda_p n_r} \mathrm{P}^+ \xrightarrow{n_p^+/\tau_p} \emptyset$$

$$\mathrm{p}^+ \underset{k_{out} n_p^-}{\overset{k_{in} n_p^+}{\rightleftarrows}} \mathrm{p}^-$$

$$\mathrm{P}^- \xrightarrow{n_p^-/\tau_p} \emptyset$$

Here $\mathrm{P}^+$ refers to the proteins in dilute phase, and $\mathrm{P}^-$ refers to the protein within the condensate. The number of mRNA, dilute phase protein and droplet phase protein molecules are denoted as $n_r$, $n_p^+$, and $n_p^-$ respectively. The volume fraction of the condensate is $\psi = n_p^- v/V_{tot}$, where $v$ is the protein molecular volume and $V_{tot}$ is the cell volume. The volume fraction (concentration) of protein in dilute phase is $\phi_+ = n_+ v/(V_{tot} - \psi V_{tot})$, and the volume fraction of protein within the condensate is $\phi_- = 1$, assuming the condensate purely consists of proteins.

System's free energy is defined as

$$F(n_+, n_-) = -\mu n_- + k_B T n_+ \left( \ln\left( \frac{n_+}{\frac{V_{tot}}{v} - n_-} \right) - 1 \right),$$

where the relative chemical potential between dilute phase and dense phase depends on the threshold hold volume fraction $\phi^*$: $\mu = -k_B T(\ln \phi^* - \phi^*)$. At equilibrium, the protein exchange rates satisfy detailed balance:

$$k_{out} = k_{in} \exp\left(-\frac{\Delta F}{k_B T}\right)$$

Rate of protein entering the condensate is assumed to be diffusion-limited, i.e., $k_{in} = 1/\tau_d = \frac{1}{(1-\psi)^{2/3}\tau_D}$, where $\tau_D$ defines the characteristic time scale of protein exchange between dilute and dense phase. A more detailed discussion about the model formulation can be found in (*1*).

Extrinsic noise of the system is modelled as the randomness in the transcription rate $\Lambda_r$, drawn from a normal distribution $N(\mu_r.\sigma_r^2)$. $\mu_r$ is selected such that the expectation of total protein volume fraction is 0.2. 100 independent and parallel simulations are performed using Gillespie algorithm(*3*), accounting for both extrinsic and intrinsic noise. The noise strength of different variables is defined as the coefficient of variance of their steady state values. The parameterization of the model is in Table S2.

**Materials and Methods**

Cell culture

*E. Coli.* BL21 (DE3) (NEB) was transformed with plasmids containing genes encoding $RLP_{WT}$ or $ELP_{VPGVG-40}$. The transformed cells were plated on a kanamycin selection plate and grown overnight. Three separate colonies were picked and cultured in 5 mL of 2x YT medium with 45 mg $mL^{-1}$ kanamycin and 1% glucose to repress basal protein expression and cultured at 37 °C at 225 r.p.m. After 16-18 h, the cells were diluted at a ratio of 1:100 into 2 x YT medium with 45 mg mL-1 kanamycin and cultured for an additional 1.5 h. The $OD_{600}$ of cells were then adjusted to 0.2 and the cells were separated into two groups as cell with condensates and without condensates. To make cells containing condensates, the culture was induced with 0.5 mM or 0.1 mM IPTG. For cells without condensates, no inducer was added. This time point was set as 0. At different time points, the cells were pelleted and washed with HBSS buffer (ThermoFisher) and the $OD_{600}$ was adjusted to 0.1 and pelleted again for further sample processing before imaging.

Confocal imaging for measuring pH

SNARF™-4F 5-(and-6)-Carboxylic Acid, Acetoxymethyl Ester, Acetate (C-SNARF4-AM, ThermoFisher) was used as the dye for pH measurement based on established protocols(*4, 5*). C-SNARF-4-AM was first dissolved in DMSO with a final concentration of 2 mM as the stock solution. The stock solution was then dissolved into HBSS buffer at desired conditions with a final concentration of 10 μM as dyeing solution. The pelleted cells were resuspended in the dyeing solution and incubated at 37 °C for 30 min and resuspended into HBSS buffer at desired conditions before imaging. The solution containing cells was put on an uncoated 35 mm Dish with No. 1.5 Coverslip with 14 mm glass (MATTEK) for imaging analysis. LEICA TCS SP8 (Leica) was used for imaging. White light laser (WLL) was used with an excitation wavelength set at 488 nm with 70 % power and 2 % laser line intensity. Hybrid detectors (HYD) were set at two different ranges for ratiometric imaging with one detector at 540-600 nm and the other at 610-660 nm. The 40x/1.25–0.75-NA oil immersion objective was used to calibrate the location and 100x/1.40 oil immersion object was used to acquire the images for single cell analysis. The acquired images were first processed in LAS X Life Science Microscope software (Leica) and quantified by ImageJ (NIH) based on the ratio between the intensity of these two channels.

Confocal imaging for measuring membrane potential

Di-4-ANEPPS (ThermoFisher) dye was used to measure membrane potential. Di-4-ANEPPS was first dissolved in DMSO with a final concentration of 1 mM as the stock solution. An imaging solution based on HBSS buffer containing 1 μM of Di-4-ANEPPS was used to resuspend the cells. The cells were then incubated at room temperature for 30 min before washing the cells with HBSS and then transferred onto an uncoated 35 mm Dish with No. 1.5 Coverslip with 14 mm glass (MATTEK) for imaging. LEICA TCS SP8 (Leica) was used for imaging. White light laser (WLL) was used with an excitation wavelength set at 498 nm with 70 % power and 10 % laser line intensity. Hybrid detector (HYD) was set at 550 nm to 680 nm. 100x/1.40 oil immersion object was used to acquire the images for single cell analysis. ImageJ was used to quantify the intensity signal at the cellular membrane.

Flow cytometry for membrane potential analysis

Thioflavin T (Millipore Sigma) was used as an indicator to measure the uptake of small molecule upon membrane hyperpolarization as a strategy to compare the change of membrane potential(*6*). Cells with or without condensates treated with different chemical clamps were incubated with 1 mM ThT at 37°C for 30 min. Cells were filtered through a 35 micron nylon filter immediately before recording the samples. BD FACSCanto-II Cytometer with a Violet laser was used to evaluate the ThT signal. Cells without incubating with ThT were used as negative controls. BD FACSDiva software was used for both data acquisition and analysis. 10,000 events were recorded for sample analysis. All events were plotted on an FSC-A vs SSC-A bivariate pseudocolor dot plot to exclude debris and to gate on healthy cells (P1 gate). Doublet discrimination was performed hierarchically on P1 population by first plotting SSC-W vs SSC-H followed by FSC-W vs FSC-H. Single cells were then analyzed for ThT level by plotting ThT area signal (log scale) against SSC-A (linear scale). Negative control was utilized to set the threshold for gating ThT+ population. Histogram of fluorescence intensity versus counts of the cells was plotted to compare the distribution of ThT signal at the population level.

<u>Inductively coupled plasma mass spectroscopy (ICP-MS) analysis of the cytoplasm solution</u>

2 L of cells were cultured in order to obtain enough cellular volume to trigger temperature dependent phase transition. The following method demonstrated the culture process of 1 L cell culture. 12 mL of cells containing the plasmid of interest was cultured overnight in 2x YT containing 45 mg $mL^{-1}$ kanamycin and 1 % glucose. Cell culture was then diluted into 1 L of 2x YT containing 45 mg $mL^{-1}$ and cultured for 3 h at 37 °C and 150 r.p.m. Cell culture was then induced with 0.5 mM IPTG and cultured for another 6 h. Cell pellets were collected by centrifuging cultures at 3,500g for 10 min at 4 °C and 2 L of cells were resuspended in 5 mL of cell lysis buffer (50 mM Tris pH 7.5), which typically resulted a total volume around 7.2 mL. Resuspended cells were lysed by 2 min of sonication (10 s of sonication followed by 40 s of rest) at an intensity of 75% in an ice bucket. This solution is the disrupted cell lysate for further processing.

For cells transformed with RLP construct, which undergoes phase transition through UCST phase behavior(*7*), the disrupted cell lysate was centrifuged at 22 °C at 20,000 g for 20 min to achieve separation of phases. For the sample at condition with condensates, the supernatant of the centrifuged solution was directly subjected to further processing. For the sample at condition without condensates, the solution with debris and condensates was put into water bath for 1 h at 37 °C to dissolve the condensates. The dissolution of the dense phase could be observed by eyes as shown previously(*7*). The supernatant was then extracted for further processing. The extracted supernatant for both samples was subjected to treatments with 1.6 U $mL^{-1}$ proteinase K (NEB) and 500 U $mL^{-1}$ Benzonase nuclease for 30 min at 37 °C before ICP-MS analysis.

For cells transformed with ELP construct, which undergoes phase transition through LCST phase behavior(*8, 9*), the disrupted cell lysate was centrifuged at 40 °C at 25,000 g for 20 min to achieve separation of phases. For the sample at condition with condensates, the supernatant of the centrifuged solution was immediately removed after centrifuge and subjected to further processing. For the sample at condition without condensates, the solution with debris and condensates was put into 4 °C to dissolve the condensates. Comparing with RLP condensates, for ELP condensates, it was difficult to observe the dissolution process by eyes because the ELP dense phase can be mixed with other cellular debris. To establish the analysis procedure, we evaluated the dissolution process by preparing multiple ELP samples and incubating the same condition prepared ELP samples at room temperature and at 4 °C. We then removed the supernatant and compared the weight of the pellet of different samples to confirm the transition process happened. The supernatant was then extracted for further processing. The extracted supernatant for both samples was subjected to treatments with 1.6 U $mL^{-1}$ proteinase K (NEB) and 500 U $mL^{-1}$ Benzonase nuclease for 30 min at 37 °C before ICP-MS analysis.

An aliquot of the supernatant solution was diluted into a 2% nitric acid (v/v) and 0.5% hydrochloric acid (v/v) solution prepared with 18.2 MΩ water and trace metal grade acids. The matrix was spiked with 72Ge and 89Y internal standards. The samples were measured for Mg, Na, and K in a collision cell mode under He gas on an Agilent 7900 ICP-MS. The instrument was calibrated with a multi-element matrix (Spex Certiprep 2A) and a second source standard was used to verify the calibration curve (CRM-TMDW-A, High Purity Standards). The data was processed based on the dilution ratio used during the process of sample processing.

<u>Simulation on stochastic gene expression</u>

Simulation of stochastic gene expression and condensate forming processes are modeled as previously reported (*1*), using Gillespie algorithm (*2*). Briefly, we consider transcription, translation, and protein exchange between dense and dilute phases:

$$\emptyset \xrightarrow{\Lambda_r} \mathrm{RNA} \xrightarrow{n_r/\tau_r} \emptyset$$

$$\emptyset \xrightarrow{\lambda_p n_r} \mathrm{P}^+ \xrightarrow{n_p^+/\tau_p} \emptyset$$

$$\mathrm{p}^+ \underset{k_{out} n_p^-}{\overset{k_{in} n_p^+}{\rightleftarrows}} \mathrm{p}^-$$

$$\mathrm{P}^- \xrightarrow{n_p^-/\tau_p} \emptyset$$

Here $\mathrm{P}^+$ refers to the proteins in dilute phase, and $\mathrm{P}^-$ refers to the protein within the condensate. The number of mRNA, dilute phase protein and droplet phase protein molecules are denoted as $n_r$, $n_p^+$, and $n_p^-$ respectively. The volume fraction of the condensate is $\psi = n_p^- v/V_{tot}$, where $v$ is the protein molecular volume and $V_{tot}$ is the cell volume. The volume fraction (concentration) of protein in dilute phase is $\phi_+ = n_+ v/(V_{tot} - \psi V_{tot})$, and the volume fraction of protein within the condensate is $\phi_- = 1$, assuming the condensate purely consists of proteins. For detailed model formulation and parameterization, see Supplementary Text 1.

To account for the heterogeneity across different cells, we considered both extrinsic and intrinsic noises. We assumed the gene expression rate of the condensate-forming protein to follow a Gaussian distribution, as the extrinsic noise, while the stochastic transcription, translation, and the protein exchange between dilute and dense phases are considered as intrinsic noise.

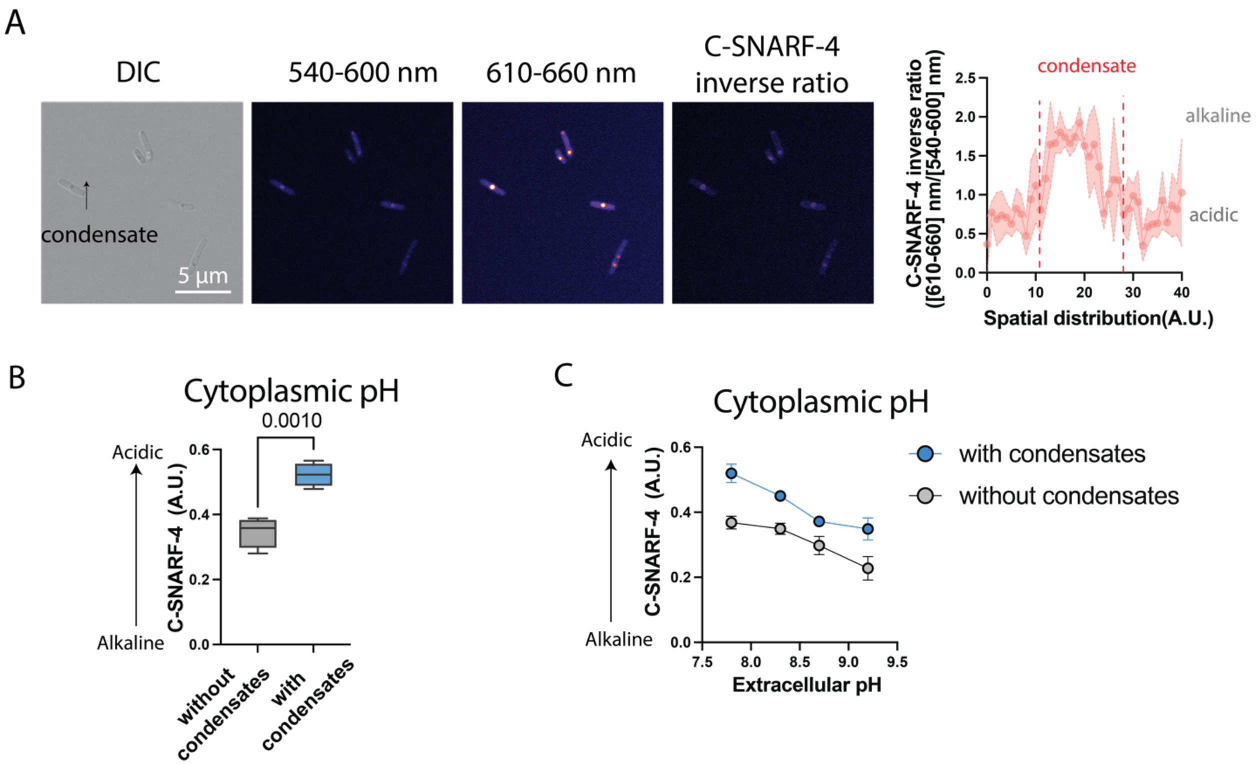


**Figure S1. Phase separation can establish a pH gradient between the cytoplasm and the condensate.**

**A)** Representative ratiometric image of a distinct pH between the dilute and the dense phases. Right panel shows the quantification of the C-SNARF-4 signal gradient across the cell.

**B)** Comparison of cytoplasmic pH based on C-SNARF-4 signal for cells without and with condensates. N = 3 independent experiments with over 30 cells quantified per experiment. Two-tailed t-test for statistical analysis.

**C)** Evaluations of the capability of condensates to modulate cytoplasmic pH under different extracellular pH conditions. N = 4 independent experiments with over 20 cells quantified per experiment.

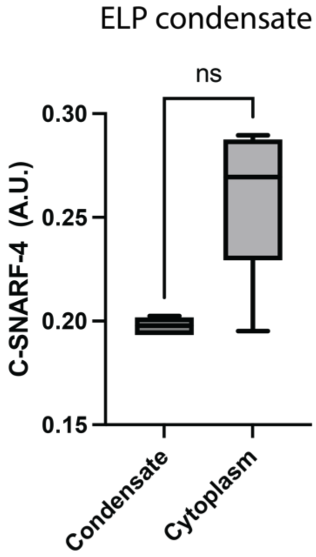


**Figure S2. Phase transition of ELP is incapable of manipulating the cytoplasmic pH.** Cells containing a plasmid encoding ELP ($[VPGVG]_{40}$) was induced with 0.1 mM IPTG. After 2 h of induction, cytoplasmic pH and condensate pH were quantified with C-SNARF-4 assay. N = 25 individual cell. Two-tailed t-test for statistical analysis. ns means non-significance.

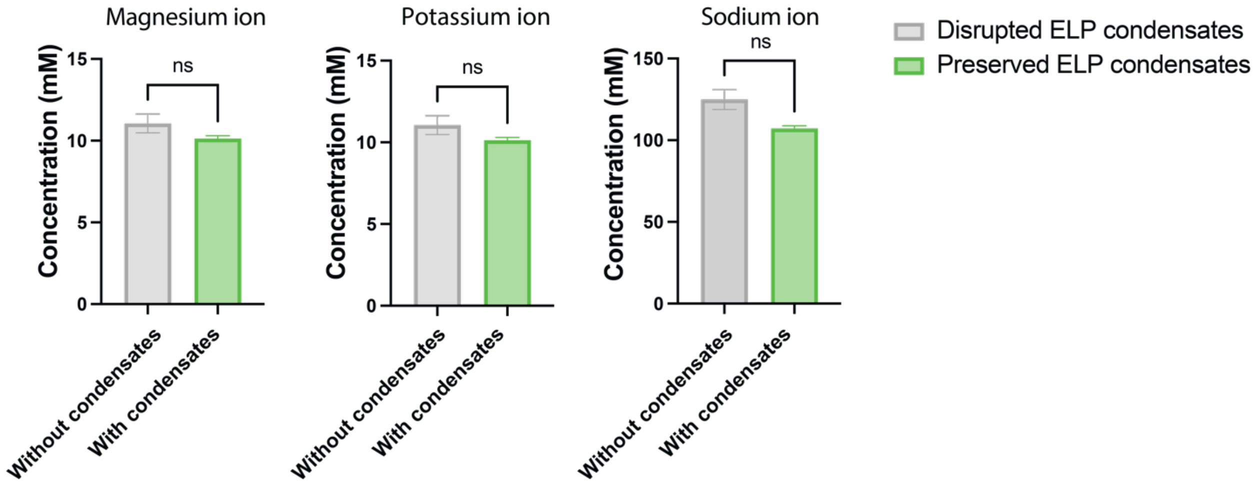


**Figure S3. Inductively coupled plasma mass spectrometry (ICP-MS) analysis of the effects of phase transition on cytoplasmic ion condition.** Cytoplasmic ion concentration of magnesium, potassium and sodium at the conditions with or without ELP condensates. Two-tailed t-test for statistical analysis. ns means non-significance.

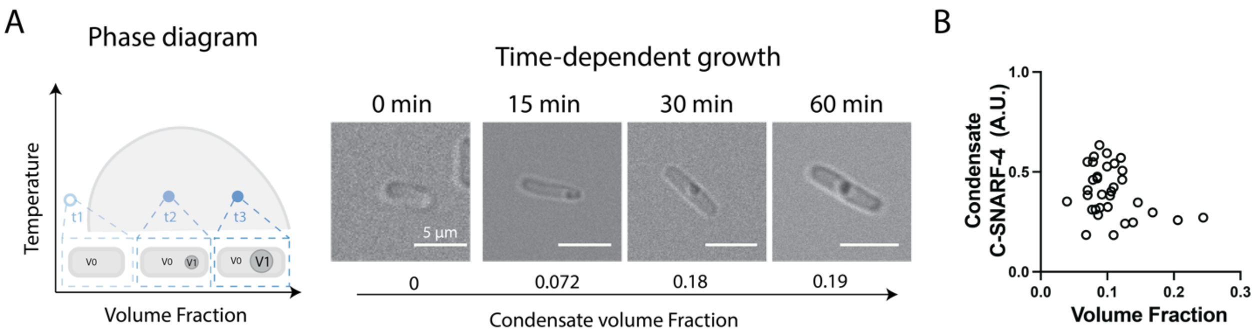


**Figure S4. Time-dependent characterization of volume fraction of condensates in a cell.**

**A)** Overexpression of RLP after forming condensates results in the increase of condensate volume fraction.

**B)** Dense phase pH is not correlated with the volume fraction of the condensates.

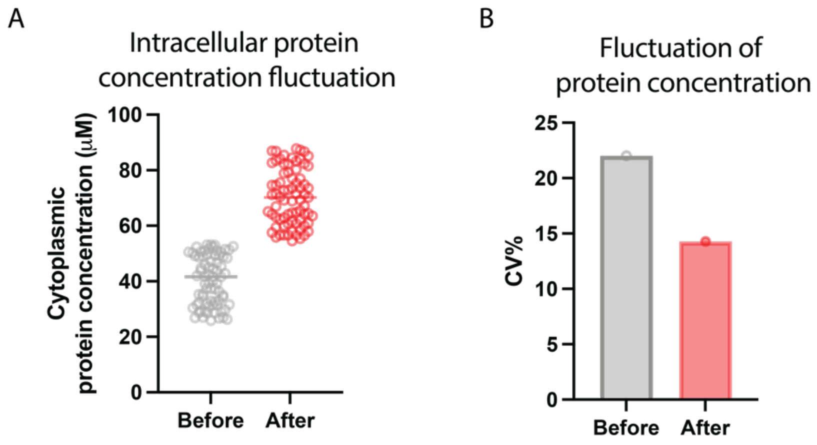


**Figure S5.** Evaluation of the cellular signal fluctuations before and after condensate formation.

**A)** Quantification of cytoplasmic RLP-mEGFP protein concentration before and after phase separation. Cells containing gene encoding RLP-mEGFP were induced with 0.05 mM IPTG. Samples without condensates were imaged after 45 min of induction. Samples with condensates were imaged after 90 min of induction. Cytoplasmic protein concentration was quantified based on previous published works(*10, 11*).

**B)** Comparison of the coefficient of variance of cytoplasmic protein concentration based on cells with and without condensates. N = 75 individual cell.

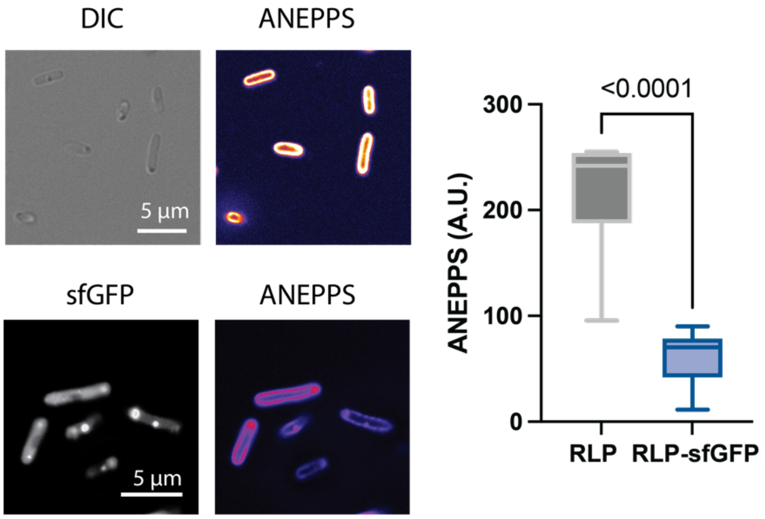


**Figure S6. Modulation of membrane potential through variation of the amount of charged residues in the protein sequence.** Cells containing genes encoding RLP or RLP-sfGFP were induced with 0.1 mM IPTG for 1.5 h before processing with ANEPPS assay. N = 25 individual cell. Two-tailed t-test for statistical analysis.

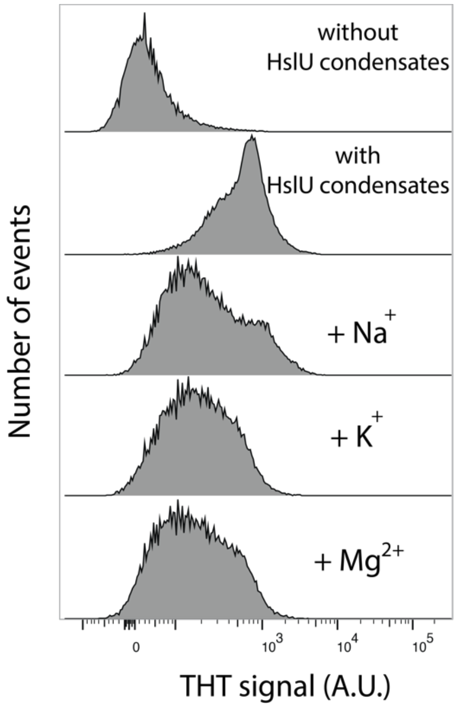


**Figure S7. Evaluation of the effect of HslU condensates on the membrane potential.** Cells containing a plasmid encoding HslU protein were induced with 0.5 mM IPTG. For cells without condensates, samples were processed with ThT assay after 3 h of induction. For cells with condensates, samples were processed with ThT assay after 7 h of induction.

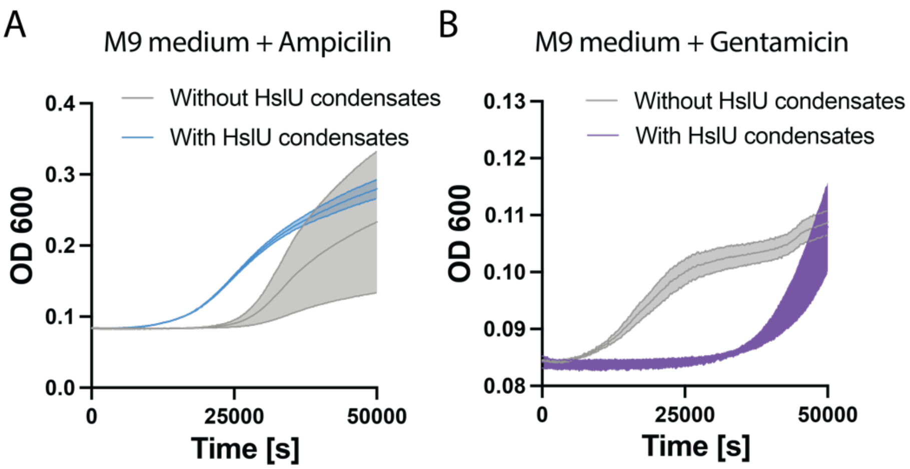


**Figure S8.** Evaluation of the effect of HslU condensates on the resistance of different antibiotics.

**A)** Evaluation of cellular growth of cells with or without HslU condensates in M9 minimum medium with 2 µg/mL ampicillin.

**B)** Evaluation of cellular growth of cells with or without condensates in M9 minimum medium with 0.5 µg/mL gentamicin.

**Table S1.** Genes used in this study.

| Plasmid | Protein sequence | Antibiotic resistance | Origin of replication |
|---|---|---|---|
| pET24+-RLP$_{WT}$ | SKGP-[GRGDSPYS]$_{20}$-GY | Kanamycin | pBR322 |
| pET24+-ELP$_{V40}$ | [VPGVG]$_{40}$-GY | Kanamycin | pBR322 |
| pET24+-RLP$_{WT}$-sfGFP | SKGP-[GRGDSPYS]$_{20}$-SKGEELFTG VVPILVELDG DVNGHKFSVR GEGEGDATNG KLTLKFICTT GKLPVPWPTL VTTLTYGVQC FSRYPDHMKR HDFFKSAMPE GYVQERTISF KDDGTYKTRA EVKFEGDTLV NRIELKGIDF KEDGNILGHK LEYNFNSHNV YITADKQKNG IKANFKIRHN VEDGSVQLAD HYQQNTPIGD GPVLLPDNHY LSTQSVLSKD PNEKRDHMVL LEFVTAAGIT HGMDELYKGY | Kanamycin | pBR322 |
| pET24+-RLP$_{WT}$-mEGFP | SKGP-[GRGDSPYS]$_{20}$- VSKGEELFT GVVPILVELD GDVNGHKFSV SGEGEGDATY GKLTLKFICT TGKLPVPWPT LVTTLTYGVQ CFSRYPDHMK QHDFFKSAMP EGYVQERTIF FKDDGNYKTR AEVKFEGDTL VNRIELKGID FKEDGNILGH KLEYNYNSHN VYIMADKQKN GIKVNFKIRH NIEDGSVQLA DHYQQNTPIG DGPVLLPDNH YLSTQSKLSK DPNEKRDHMV LLEFVTAAGI TLGMDELYKGY | Kanamycin | pBR322 |
| pET24+-HslU | MSEMTPREIVSELDKHIIGQDNAKRSVAIALRNRWRRMQLNEELRHEVTPKNILMIGPTG VGKTEIARRLAKLANAPFIKVEATKFTEVGYVGKEVDSIIRDLTDAAVKMVRVQAIEKNR YRAEELAEERILDVLIPPAKNNWGQTEQQQEPSAARQAFRKKLREGQLDDKEIEIDLAAA PMGVEIMAPPGMEEMTSQLQSMFQNLGGQKQKARKLKIKDAMKLLIEEEAAKLVNPEELK QDAIDAVEQHGIVFIDEIDKICKRGESSGPDVSREGVQRDLLPLVEGCTVSTKHGMVKTD HILFIASGAFQIAKPSDLIPELQGRLPIRVELQALTTSDFERILTEPNASITVQYKALMA TEGVNIEFTDSGIKRIAEAAWQVNESTENIGARRLHTVLERLMEEISYDASDLSGQNITIDADYVSKHLDALVADEDLSRFIL | Kanamycin | pBR322 |

**Table S2**. Parameters used for simulating condensate formation process.

| $\mu_r$ | $\sigma_r/\mu_r$ | $\lambda_p$ | $\tau_r$ | $\tau_p$ | $\tau_D/\tau_p$ | $V_{tot}/v$ | $\phi^*$ |
|---|---|---|---|---|---|---|---|
| $0.11\ s^{-1}$ | $0.011 s^{-1}$ | $0.005\ s^{-1}$ | $100\ s$ | $3.6\times10^3\ s$ | 0.002 | 1000 | 0.1 |
